# Reelin haploinsufficiency affects skilled motor performance associated with suppression of training-induced gene enrichment, synaptic function and activity-dependent cortical plasticity in mice

**DOI:** 10.1101/2020.10.25.351528

**Authors:** Mariko Nishibe, Hiroki Toyoda, Yu Katsuyama

**Affiliations:** Office of Strategic Innovative Dentistry, Graduate School of Dentistry, Osaka University, Osaka, 565-0871 Japan; Department of Anatomy, Shiga University of Medical Science, Shiga, 520-2192 Japan; Department of Oral Physiology, Graduate School of Dentistry, Osaka University, Osaka, 565-0871 Japan

## Abstract

RELN (Reelin) is one of the genes implicated in neurodevelopmental psychiatric vulnerability. Patients with neurodevelopmental disorders can experience impairments in fine motor skills. While Reelin modulates synaptic function, whether Reelin haploinsufficiency affects activity-dependent cortical plasticity which supports development of skilled movement is unclear. Here, heterozygous *Reeler* mutant (HRM) and *Dab1* ^*floxed*/ +^; *Emx1-Cre* mice both displayed learning improvements measured by the reach-to-grasp task, but their performance levels of the forelimb motor skill were lower, compared with controls. The level of skilled motor performance was correlated with the area of cortical representations of the trained forelimb, examined after 10 days of training. Furthermore, we hypothesized that the genetic haploinsufficiency also alters changes that occur during the early phase of the training. Examined on day 3, the training induced synaptic modifications of the layer III cortical neurons in (wild-type) WT mice, which were contributed by synaptic potentiation and increase in spontaneous action-potential driven glutamatergic-transmission. On the other hand, the basal excitatory and inhibitory synaptic function were depressed, affected both by presynaptic and postsynaptic synaptic impairments in naive HRM; and thus, no further training-induced synaptic plasticity occurred in HRM. Lastly, examined after 3 days of training, the gene enrichment observed in trained WT mice was absent in trained HRM mice. The finding suggests the Reelin haploinsufficiency alters the skilled motor function; and we propose the suppression of gene enrichment, and synaptic abnormality led by the genetic insufficiency may contribute to impede the occurrence of activity-dependent cortical plasticity.

**Significance Statement:** Impairments in fine motor skills occur in subjects with neurodevelopmental disorders. We report a mutation relevant to the neurodevelopmental disorders can impact the cortical plasticity associated with skilled motor function. In wild-type mice, the motor training induced extensive activity-dependent cortical map plasticity, synaptic modifications through synaptic potentiation and excitatory-transmission increase, as well as enrichments in certain gene expressions. On the other hand, mice with Reelin haploinsufficiency (presumed mouse model of neurodevelopmental disorders) exhibited lower level of skilled motor performance, and the underlying correlates shown in wild-type mice were found suppressed. We conclude the suppression of gene enrichment, and synaptic abnormality due to Reelin haploinsufficiency may underlie the limited development of activity-dependent cortical plasticity, contributing to impairments in motor skills.

## Introduction

The patients with neurodevelopmental psychiatric vulnerability often experience impairments in fine motor skills. The reach and grasp movements are altered in children with autism spectrum disorders (Mari et al., 2003), which can be detected as early as age 3 (Sacrey et al., 2018). Several fMRI studies have identified that fine motor skill impairments in subjects with schizophrenia correlate with the hypofunction of the primary motor cortex (M1) (Schroder et al., 1995; Thompson et al., 2017). It has been shown that topographical map reorganization of the motor cortex, through changes in the horizontal cortical connectivity and redistribution of movement representation, is one mechanism by which mammals acquire skilled movements (Hess and Donoghue, 1994; Castro-Alamancos et al., 1995; Kleim et al., 1998; Rioult-Pedotti et al., 1998).

Genetic association studies on neurodevelopmental diseases find causative involvement of a number of genes, including *RELN* (Reelin). And, another causative gene strongly implicated in Schizophrenia, disrupted-in-schizophrenia (*DISC1*) (Facal and Costas, 2019), is linked to the regulation of synaptic development and to the processing of Reelin (Bradshaw et al., 2020). Reelin is an extracellular glycoprotein highly conserved in the chordates. Upon Reelin binding to its receptors, DAB Adaptor Protein 1 (Dab1) becomes phosphorylated by Src family kinases, thereby, activating intracellular mechanisms affecting synapses. Specifically, Reelin has been demonstrated to modulate the molecular composition of the synapses (Chen et al., 2005; Sinagra et al., 2005; Groc et al., 2007; Ventruti et al., 2011), and to regulate the functions of presynaptic structural organization (Hellwig et al., 2011) and postsynaptic dendritic spine formation (Pappas et al., 2001; Niu et al., 2004; Pujadas et al., 2010). In the heterozygous *reeler* mutant (HRM) mice, several behavioral impairments have been reported including deficits in prepulse inhibition (PPI), hippocampus-dependent memory (Weeber et al., 2002; Qiu et al., 2006; Ognibene et al., 2007; Rogers et al., 2013), and in the sensorimotor gating (Tueting et al., 1999; Barr et al., 2008; Teixeira et al., 2011). Further, the *in vivo* supplementation of Reelin or, *in vivo* application of NMDA receptor blockers has been demonstrated to augment hippocampal LTP and to rescue behavioral impairments related to hippocampus function in HRM (Rogers et al., 2013; Iafrati et al., 2014).

It is unknown, however, whether Reelin haploinsufficiency affects fine motor skill and activity-dependent cortical plasticity. Here, we found HRM and Dab1 floxed/ +; Emx1-Cre (*Dab1* cKO) mice showed motor learning improvements while subjected to the single pellet reach-to-grasp training, but they displayed impaired level of skilled motor performance. Their behavioral development was accompanied by less extensive map enlargement representing the movements of the trained forelimb in M1 after 10 days of training, assessed by intracortical microstimulation. Previous studies have shown the map reorganization induced by motor skill training occurs during the late phase of the training representing the consolidation of the skill (Kleim et al., 2004), in contrast to rapid improvements in behavior, and dendritic pruning and hypertrophy detected early in the training (Greenough et al., 1985; Xu et al., 2009). Hence, the present study further examined the effect of Reelin haploinsufficiency on changes that may precede the map plasticity. In WT mice after 3 days of training, we observed an increase in the spontaneous action potential driven glutamatergic transmission, thereby providing a putative mechanism for enhanced LTP expression at the M1 excitatory synapses. In HRM, such changes did not occur by training, due to abnormality in the synaptic formation and function in the naïve HRM. Specifically, the whole-cell recording showed lower basal LTP, and lower basal inhibitory and excitatory synaptic transmissions, suggesting that the activity-dependent cortical plasticity requires the basal synaptic function of normal development. Lastly, RNA-sequencing analysis suggests HRM exhibited substantially lower gene enrichment in response to 3 days of training than WT. We propose that the abnormality led by the genetic insufficiency may be closely linked to the suppression of activity-dependent cortical plasticity, at least partially underlying the impairments in the skilled motor function.

## Materials and Methods

Animal use was approved by the Guidelines for the Care and Use of Laboratory Animals both at Shiga University of Medical Science and of the Graduate School of Dentistry at Osaka University. All efforts were taken to minimize suffering. We used a total of 125 {53 WT mice, 52 heterozygous *reeler* mice (*rl*/ +) of B6C3Fe-*a/a-Relnrl* strain (RBRC00045, Riken BRC, Tsukuba Japan); and 10 *Dab1* ^*floxed*/ +^; *Emx1-Cre*, and 10 *Dab1* ^+/+^; *Emx1-Cre* mice (Imai et al., 2017)} of either sex randomly assigned. Sex distribution in each experiment was made as equal as possible while mice from the same litters were used. Sex distribution is provided in each experiment description. The age of the mice used for the patch clamp was 5 weeks (10 WT and 10 HRM mice) and for the rest of all experiments 17-20 weeks (43 WT, 42 HRM, 10 *Dab1* cKO, 10 *Dab1* control mice). Genotyping was conducted as previously described (Nishibe et al., 2018). 3-4 mice were housed with ad libitum food (except the time of food regulation during the training, described below) and water on a 12h: 12h light: dark cycle, with ambient temperature maintained at 24°C.

### Reach-to-grasp training

Pellet retrieval task (Whishaw et al., 1992) was used to train the forelimb motor skills in mice. The testing box was made of Plexiglas, measured 13 cm (length) * 9cm (width) * 15 cm (height). A 1 cm wide reaching slot, extending from the bottom of the reaching box to 7 cm in height, was centered at the front of the chamber. A food shelf was secured to the outside of the chamber 1.5 cm from the bottom. The behavioral study (Figure 1A) included WT and HRM (n = 10, Movie 1 and Movie 2; WT and HRM group each consisted of 6 male mice and 4 female mice), and *Dab1* control and *Dab1* cKO mice (n = 5, Movie 3 and Movie 4; the two groups each consisted of 2 male and 3 female mice). For the rest of the experiments, all mice were either assigned to undergo reach to grasp training or assigned to eat pellets from the floor of the reaching box, both followed by the acclimation and behavioral shaping procedure with similar amount of handling by the experimenters. The mice were sacrificed on 3 days of training for the patch-clamp, qRT-PCR, and RNA-sequencing sample preparation, and after 10 days of training for the ICMS experiment.

**Figure 1.**
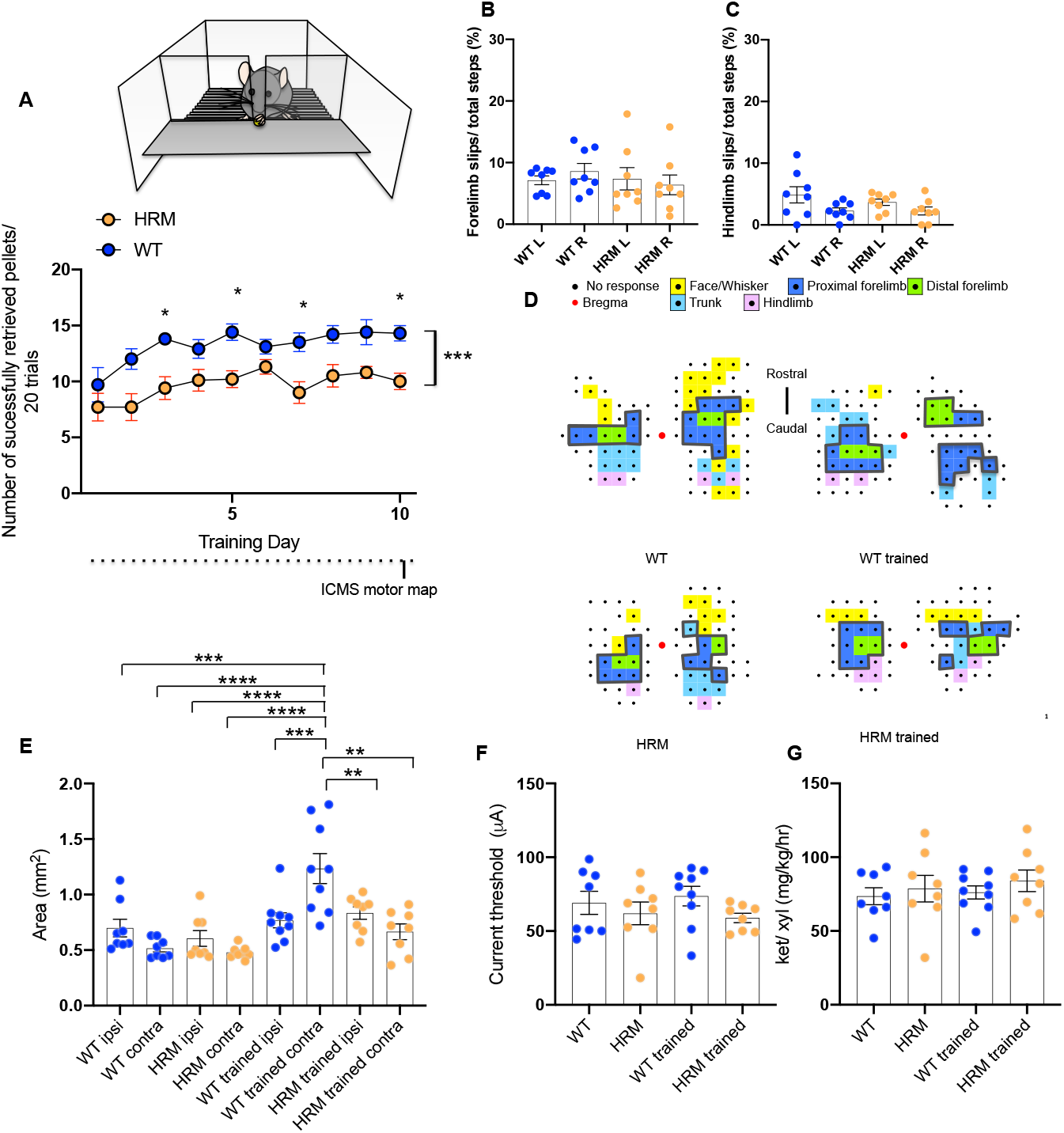
Skilled motor behavior and M1 forelimb map of WT and HRM. **(A)** TOP: schematic drawing of pellet reach-to-grasp task. BOTTOM: learning curve of WT and HRM mice trained at reach-to-grasp task. While both groups showed incremental improvements over 10 days, the HRM mice retrieved a fewer number of pellets per 20 trials than the WT (n = 10 mice), specifically on day 3, 5, 7, and day 10 (** p < 0.01 two-way repeated-measures ANOVA significant Group effect, *p < 0.05 by Sidak’s multiple comparison test). **(B)** Average percent foot-slips by left (L) and right (R) forelimb of WT and HRM during the grid walk test (n = 8 mice) **(C)** Average percent foot-slips by left (L) and right (R) hindlimb of WT and HRM during the grid walk test (n = 8 mice). **(D)** Representative M1 map of each group, derived after the 10-day training, by ICMS. Each dot in the map indicates a stimulation site (250 μm apart from one another). The forelimb areal measurements of M1 are the total area lined by gray line combining the proximal forelimb area designated by blue, and the distal forelimb area designated by green. In the trained groups, the map area contralateral side to the trained forelimb is shown on the right. **(E)** Total area of the forelimb representation in WT, HRM. WT trained and HRM trained. Area enlargement in forelimb representation was observed in WT trained mice. (n = 8 Tukey post-hoc test **p < 0.01, ***p < 0.001, ****p < 0.0001). The *ipsi* and *contra* labels on the x-axis indicate ipsilateral and contralateral motor cortical area to the trained forelimb, respectively. In WT and HRM, ipsi is the left cortex representation and contra is the right cortex representation. **(F)** The average ICMS current threshold of WT, HRM, WT trained and HRM trained group (n = 8). We found that neither Reelin haploinsufficiency nor the reach-to-grasp training affected the cortical stimulation threshold to evoke forelimb movements. **(G)** Average anesthetics volume used during the ICMS in WT WT, HRM, WT trained and HRM trained group (n = 8). The volume of anesthetics mice received during the mapping procedure did not differ among the four groups Data are presented as mean ± SEM.

#### Acclimation and behavioral shaping

Each mouse was acclimated to the human handling, and to the training and testing environment. Shaping consisted of having mice reach out for a singly-presented food pellet, about 0.5 cm from the front of the chamber. Forelimb dominance was evaluated during 20 tallied reaches, and dominance was defined in each mouse as the limb used on more than 70% of the reaches. Mice food intake was moderately regulated to 1.3-1.6 grams rodent chow per day per animal, to increase their motivation for retrieving pellets.

#### Reach-to-grasp training

Once forelimb dominance was established, we began each training session, which consisted of 20 trials. If a mouse did not complete the 20 trials within 20 minutes, one session was divided by a break, until each mouse completed the 20 trials. Thus, mice consumed at a minimum of 20 pellets of 20mg during the training. Training sessions were conducted 5 days per week for 2 weeks. A trial was counted as successful when the mouse grasped and transported the pellet to the mouth. A failure was tallied when 1) a pellet was successfully grasped, but dropped before reaching the mouth, 2) a pellet was dislodged from the food shelf, or 3) the mouse failed to contact the pellet after 5 reaches (e.g., mouse grasping air without contacting with the pellet). Trials were not tallied when a pellet was retrieved with an aid of forceps, or when a retrieval involved contact with any part of the tongue/ mouth or the non-dominant forepaw. Mice in the no training control group ate the pellets made available on the floor of the reaching box.

### Grid-walk test

To assess forelimb and hindlimb performance levels during locomotion, a grid walk task was conducted on the same group of animals used for the reach-to-grasp behavioral assessment (Figure 1B, C n = 8, WT and HRM group each consisted of 4 male mice and 4 female mice). Each mouse was placed onto an elevated grid (25cm * 35cm with 2cm * 2cm grid openings) and allowed to walk freely for 4 minutes. A foot-slip was defined as extension of the forepaw or the hindpaw through the grid opening without any of the digits catching on the grid. The number of steps and the number of slips made by each forelimb and hindlimb were recorded. Performance level was calculated as the percentage of foot-slips per step taken.

### Intracortical microstimulation (ICMS)

Upon completion of 10 days of training, each mouse was subjected to motor cortical mapping (Nishibe et al., 2018). WT group consisted of 4 male mice and 4 female mice (n = 8-9). HRM group consisted of 3 male and 5 female mice. WT trained group consisted of 4 male and 5 female mice. HRM trained group consisted of 3 male and 5 female mice. The *Dab1* cKO group consisted of 2 male and 1 female mice (n = 3-4). *Dab1* control group consisted of 1 male and 2 female mice. *Dab1* control trained group consisted of 2 male and 1 female mice. *Dab1* cKO trained group consisted of 2 male and 2 female mice. Each mouse was anesthetized with an initial dose of ketamine (100 mg/ kg IP) and xylazine (5 mg/ kg IP). Additional doses of ketamine (20 mg/ kg/ hr IM) were used when necessary to maintain adequate anesthesia (Figure 1G and Figure 2E). A craniectomy was made over the frontal cortex (2.0 mm anterior to 2.0 mm posterior to the bregma, and 0.5 to 3.0 mm lateral to the bregma). A high-magnification digital image containing a micro-scale (3R Anyty camera) of the cortical surface vasculature was obtained and imported to Photoshop. Grid lines (250*250 μm^2^) were overlaid on the digital photograph to reference the placement of the microelectrode. The bipolar electrode (FHC; Bowdoin, ME, USA) was positioned sequentially at each of the grid intersections and lowered perpendicular to the cortical surface using a hydraulic micromanipulator (Narishige MO-10, Tokyo, Japan). ICMS pulses consisted of a train of 13, 200 μs monophasic cathodal pulses at a rate of 350 Hz delivered at the rate of 1/ sec (Multi Channel System MCS GmbH, STG 4002, Reutlingen, Germany). Movements elicited by ICMS stimulation were inspected visually and recorded. The areal size was measured from the reconstructed movement maps using NIH IMAGEJ 1.50a. The distal forelimb area, as a definition, included the digits and wrist movement representations. The proximal forelimb area included the area that evoked elbow movements. At the end of the mapping procedure, each mouse was euthanized by an overdose of anesthesia cocktail of 7.5 μg medetomidine, 40 μg midazolam, and 50 μg butorphanol in 100 μL, and perfused with a transcardiac method with 0.1M PB.

**Figure 2.**
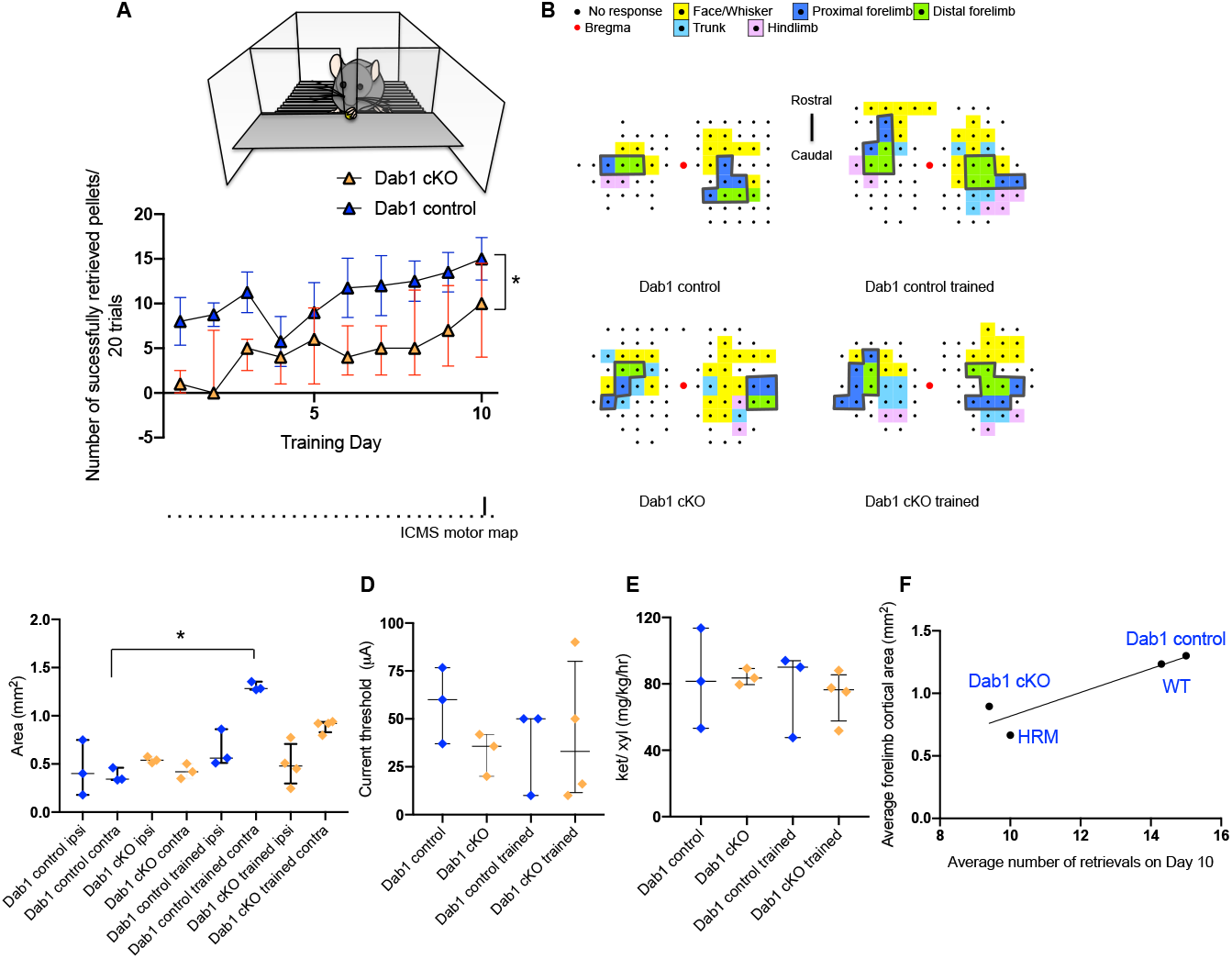
Behavior and ICMS M1 forelimb area of *Dab1* control and *Dab1* cKO mice. **(A)** TOP: schematic drawing (B) of reach-to-grasp learning task. BOTTOM: learning curve of *Dab1* ^*floxed*/ +^; *Emx1-Cre* (*Dab1*) cKO mice and *Dab1* control mice trained at reach-to-grasp task (n = 5). While both groups showed an increase in the level of performance, *Dab1* cKO mice retrieved a significantly lower number of pellets in 20 trials (p* < 0.05 two-way repeated-measures ANOVA significant Group effect). Data are presented as mean ± SEM. **(B)** Representative cortical motor map of the groups, each derived by ICMS. **(C)** Scatter plots of the total forelimb area of M1 (n = 3-4). A unilateral expansion in the size of the forelimb area, contralateral to the trained forelimb was observed in the *Dab1* control trained group (*p < 0.05 Dunn’s post hoc test). **(D)** Average ICMS current threshold of *Dab1* control and *Dab1* cKO group with and without the training (n = 3-4 mice). No effect of Dab1 haploinsufficiency or a training effect was observed on the cortical stimulation threshold. **(E)** Average volume of anesthetics used during the ICMS in *Dab1* control and *Dab1* cKO group with and without the training (n = 3-4 mice). The volume of anesthetics each mouse received was ensured not significant. Data are presented as median ± interpercentile range. **(F)** The larger the cortical map representing the trained forelimb, the better the behavioral performance level displayed in the mouse models of neurodevelopmental disorder on training day 10 (Pearson correlation coefficient p < 0.05).

### Whole-cell patch-clamp recording

The whole-cell patch-clamp recordings on neurons in 5 week-old mice produced sharp readings *in vitro* slice preparation. WT group consisted of 4 male and 1 female mice (n = 5). HRM group consisted of 3 male and 2 female. WT trained group consisted of 2 male and 3 female. HRM trained group consisted of 4 male and 1 female mice. These HRM mice exhibited a lower performance level in the pellet reach to grasp task, similar to the mice in ICMS experiment (Figure 3A). Within 30 minutes of the completion of day 3 training, mice were anesthetized with isoflurane, and the brain was quickly removed from the skull and immersed in ice-cold modified artificial cerebrospinal fluid (aCSF) composed of 210 mM sucrose, 2.5 mM KCl, 2.5 mM MgSO4, 1.25 mM NaH2PO4, 26 mM NaHCO3, 0.5 mM CaCl2 and 50 mM d-glucose. With a microslicer (Linearslicer Pro 7, Dosaka EM, Kyoto, Japan), coronal sections of 300 μm thickness were prepared. Slices were incubated at 32 °C for 30 min in 50% modified aCSF and 50% normal aCSF (pH 7.3) composed of 126 mM NaCl, 3 mM KCl, 1 mM MgSO4, 1.25 mM NaH2PO4, 26 mM NaHCO3, 2 mM CaCl2, and 10 mM d-glucose. Using MultiClamp 700B Amplifier (Molecular Devices, Foster City, CA), whole-cell recordings were made from visually identified pyramidal neurons in layer III of the motor cortex, visualized using differential interference contrast microscopy (BX-51WI; Olympus, Tokyo), within the range of 0.5 mm anterior to 0.5 mm posterior to the bregma, and the 1.0 mm to 2.0 mm lateral from the midline. For excitatory postsynaptic currents (EPSCs), recording pipettes (3–5 MΩ) were filled with solution containing 123 mM K-gluconate, 18 mM KCl, 14 mM NaCl, 2 mM ATP-Mg, 0.3 mM GTP-Na3, 10 mM 4-(2-hydroxyethyl)-1-piperazineethanesulfonic acid (HEPES), and 0.2 mM ethylene glycol tetraacetic acid (EGTA); pH 7.3, adjusted with KOH. EPSCs were recorded at a holding potential of −70 mV, by repetitive stimuli (duration is 100 μs, intensity is adjusted to induce EPSCs with an amplitude of 50–100 pA) at 0.033 Hz in the presence of 10 μM bicuculline (Matsuyama et al., 2008). For inhibitory postsynaptic currents (IPSCs), the recording pipettes (3–5 MΩ) were filled with solution containing 130 mM Cs-gluconate, 10 mM CsCl, 2 mM MgCl2, 2 mM ATP-Na2, 0.4 mM GTP-Na3, 10 mM HEPES, and 0.2 mM EGTA; pH 7.3, adjusted with CsOH (Toyoda et al., 2015). The Cl- equilibrium potential was calculated to be –57 mV. IPSCs were recorded at a holding potential of 0 mV. The IPSC recordings started 10 min after establishing the whole-cell configuration and clamping at 0 mV, in the presence of 10 μM DNQX (6, 7-dinitroquinoxaline-2, 3-Dione) and 50 μM AP-5 (dl-2-amino-5-phosphonopentanoic acid). The induction protocol for LTP, involved presynaptic 80 pulses at 2 Hz with postsynaptic paring at +30 mV (Zhao et al., 2005). Series resistance was kept at 15-25MΩ and data were discarded if access resistance changed more than 10% during an experiment.

**Figure 3.**
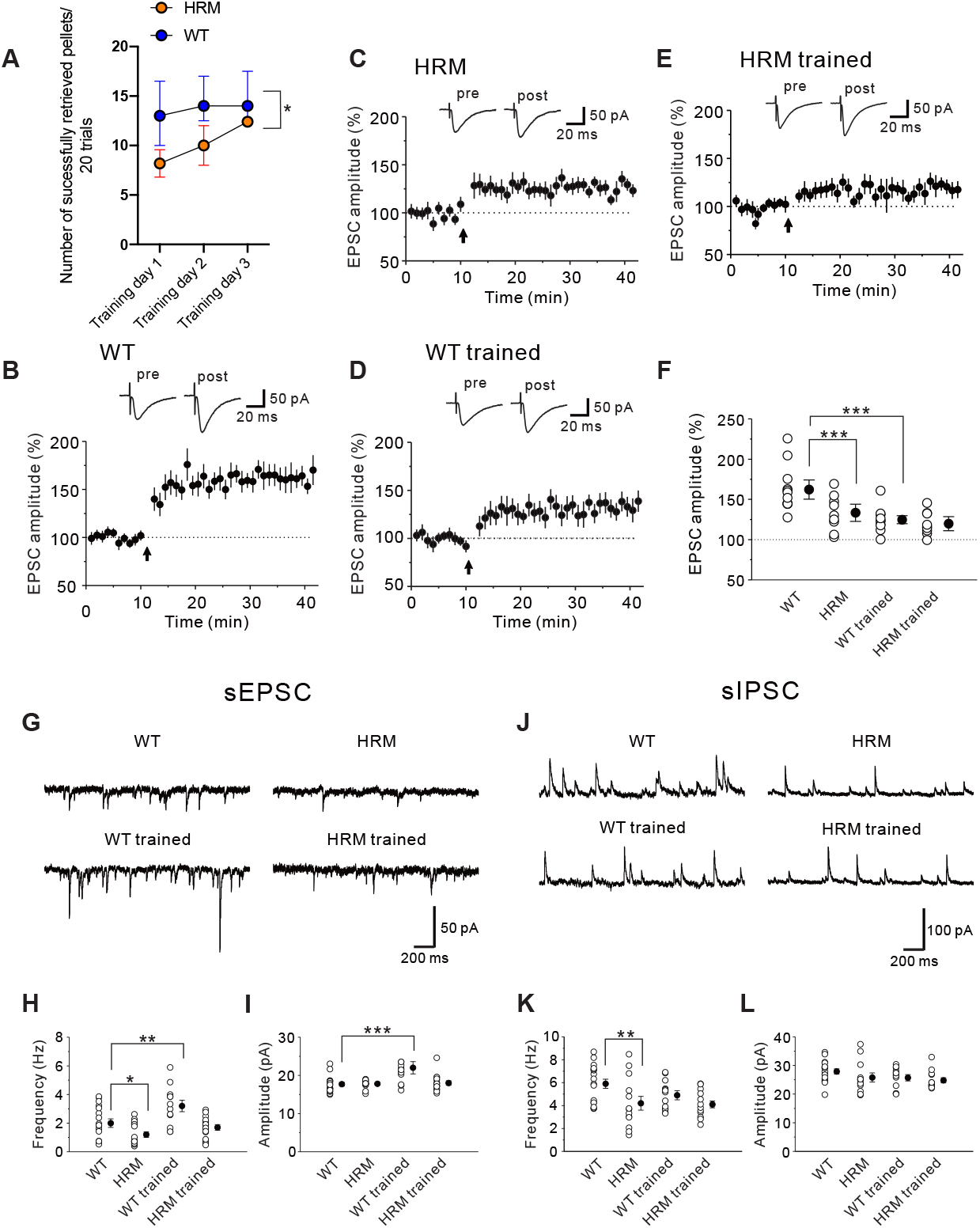
Long-term potentiation and synaptic transmissions in WT and HRM. **(A)** Behavior of the mice used in patch-clamp experiments. The HRM retrieved a fewer pellets than the WT (n = 5 mice, * p < 0.05 two-way repeated-measures ANOVA significant Group effect), similar to the mice used in rest of the study. The patch-clamp experiment proceeded on the 3^rd^ training day upon completion of the training session. **(B)** Representative insets showing averages of ten consecutive current traces before (pre) and 25-30 min after the paring (post), indicated by an arrow. The dashed line indicates the mean basal synaptic response. Synaptic potentiation, measured in excitatory postsynaptic currents (EPSC), was induced in layer III pyramidal neurons of the motor cortex in WT mice (n = 12 neurons/ 5 mice, p < 0.001, paired t-test 2-tailed), **(C)** in HRM mice (n = 9 neurons/ 5 mice, p < 0.001, paired t-test 2-tailed), **(D)** in WT trained mice (n = 10 neurons/ 5 mice, p < 0.001, paired t-test 2-tailed), and **(E)** in HRM trained mice (n = 9 neurons/ 5 mice, p < 0.001, paired t-test 2-tailed). **(F)** Summary scatter plots and average of the normalized EPSC amplitude at respective conditions (***p < 0.001, one-way ANOVA LSD post-hoc test). The dashed line indicates the mean basal synaptic response. **(G)** Representative traces of sEPSCs recorded from cortical pyramidal neurons at respective conditions. Data were collected from WT n = 18 neurons/ 5 mice; HRM n = 16 neurons/ 5 mice; WT trained n = 12 neurons/ 5 mice; and HRM trained n = 16 neurons/ 5 mice. **(H)** Summary scatter plots and average frequency of sEPSCs (*p < 0.05, **p < 0.01, one-way ANOVA LSD post-hoc test). **(I)** Summary scatter plots and average amplitude of sEPSCs (***p < 0.001, one-way ANOVA LSD post-hoc test). **(J)** Representative traces of sIPSCs recorded from cortical pyramidal neurons at respective conditions. Data were collected from WT n = 17 neurons/ 5 mice; HRM n = 13 neurons/ 5 mice; WT trained n = 13 neurons/ 5 mice; and HRM trained n = 17 neurons/ 5 mice. **(K)** Summary scatter plots and average frequency of sIPSCs (**p < 0.01, one-way ANOVA LSD post-hoc test). **(L)** Summary scatter plots and average amplitude of sIPSCs. Data are presented as mean ± SEM.

### Quantitative Reverse Transcription Polymerase Chain Reaction (qRT-PCR)

Within 30 minutes of the completion of training day 3 reach-to-grasp, mice were perfused with 0.1 M PB, under influence of overdosed anesthesia cocktail of 7.5 μg medetomidine, 40 μg midazolam, and 50 μg butorphanol in 100 μL. For the qRT-PCR results in the Figure 5, each 4 group included 6 male and 2 female mice (n = 8). For the qRT-PCR results in the Figure 6, each 4 group included 4 male and 2 female mouse samples from the mice used in the experiment in Figure 5 (n = 6). The sensorimotor cortex (0.4 mm anterior and 0.4 mm posterior from the bregma, and 1.0 mm lateral to 1.8 mm lateral from the midline) was collected from the right hemisphere of the untrained HRM and WT mice, and from the contralateral hemisphere of the trained forelimb from the trained HRM and WT mice. Total RNA was isolated using Trizol reagent (Invitrogen, CA, USA), followed by reverse transcription using the High Capacity cDNA Reverse Transcription Kit (Applied Biosystems, CA, USA). Quantitative RT-PCR was performed using Fast SYBR Green Master Mix (Applied Biosystems) and QuantStudio 7 Flex Real-Time PCR System. cDNA samples corresponding to the final RNA amount of 6-9 ng were applied to the MicroAmp Fast Optical Well Reaction Plate. The PCR condition was as follows: 95 °C for 20 sec followed by 40 cycles of 95 °C for 3 min, and 60 °C for 30 sec. The primers used are listed in Extended Data Figure 5-1 and Figure 6-1. The expression levels were calculated by the relative standard curve method using a reverse-transcribed product from the WT untrained group as standards and normalized by the geometric mean of the expression levels of Actb and ribosomal protein S18 (setting the value in the WT untrained group as 1).

### RNA-sequencing

Total RNA isolated using Trizol reagent (Invitrogen) for the qRT-PCR experiment was used for the RNA-sequencing (n = 3). WT group consisted of 1 male and 2 female mice. HRM consisted of 2 male and 1 female mice. WT trained group consisted of 2 male and 1 female mice. HRM trained group consisted of 2 male and 1 female mice. Each cDNA was generated using the High Capacity cDNA Reverse Transcription Kit (Applied Biosystems). Each library was prepared using a Nextera XT DNA Library Prep Kit (Illumina, San Diego, CA, USA) according to the manufacturer’s instructions. Whole transcriptome sequencing was applied to the RNA samples with the use of an Illumina HiSeq 2500 platform in a 75-base single-end mode. Sequenced reads were mapped to the mouse reference genome sequences (mm10) using TopHat ver. 2.0.13 in combination with Bowtie2 ver. 2.2.3 and SAMtools ver. 1.0. The number of fragments per kilobase of exon per million mapped fragments (FPKMs) was calculated using Cuffnorm ver. 2.2.1. The DEGs and pathway analysis were performed on iDEP.90 (Ge et al., 2018). For DEGs analysis all group comparisons false discovery rate (FDR) cutoff value of 0.15001, and minimum fold change value of 2 were used. Pathway significance cutoff (FDR) was 0.1, and any genes >FDR 2 was dropped prior to pathway analysis. The RNA-sequencing datasets are deposited in DNA DataBank of Japan https://www.ddbj.nig.ac.jp/index-e.html., which can be retrieved by the Experiment Accession of E-GEAD-339, and Access key of uoWCIcyXCh9qP2hDMH1D.

### Statistical Analysis

Normal distribution of samples was assessed by Shapiro-Wilkinson normality test. Normal distribution was tested using two-tailed t-test, one-way ANOVA or two-way repeated measures ANOVA followed by Sidak’s, Tukey’s, or LSD multiple comparison tests when appropriate. When data distribution was not normal we used Kruskal Wallis tests followed by Dunn’s multiple comparison tests when appropriate. Statistical analyses were performed using Prism 8.4.3 (Graphpad, CA USA). Data analyzed using parametric analysis and data analyzed by non-parametric analysis are presented as mean ± SEM and median ± interquartile range, respectively. Each statistical test used per analysis ran is indicated in the Results section. A p-value < 0.05 was considered statistically significant.

## Results

### Heterozygous *reeler* mutant mice and in *Dab1* cKO mice showed impairments in skilled motor task, accompanied by less extensive activity-dependent cortical plasticity than controls

WT mice and HRM mice were trained to reach with a preferred forepaw and to retrieve a singly presented food pellet using grasping and pronating movements (**Movie 1 and Movie 2**). While both groups showed incremental improvements over 10 days, the performance levels exhibited by HRM were lower compared with these exhibited by WT throughout the training period (**Figure 1A** n = 10 mice, group effect F_1, 18_= 19.05, p = 0.0004; time effect F_3.485, 62.73_ = 5.473, p = 0.0013; interaction effect F_9, 162_ = 0.8954, p = 0.530 two-way repeated-measures ANOVA). The WT retrieved a higher number of pellets per 20 than HRM statistically on day 3 (WT 13.8 ± 0.44 pellets, HRM 9.4 ±1.02, p = 0.018, Sidak’s multiple comparisons test), day 5 (WT 14.4 ± 0.748, HRM 10.2 ± 0.74, p = 0.0095), day 7 (WT 13.5 ± 0.83, HRM 9.0 ± 0.97, p = 0.025), and day 10 (WT 14.3 ± 0.68, HRM 10.0 ± 0.74, p = 0.0049). Meanwhile, the HRM showed intact gross motor function, tested on the grid-walk test in administered one assessment day (**Figure 1B, C** n = 8, percent foot-slips of WT left forelimb 7.1 ± 0.71%, WT right forelimb 8.5 ± 1.26%, HRM left forelimb 7.3 ± 1.80%, HRM right forelimb 6.37 ± 1.60%; F _3, 28_ = 0.427, p = 0.734 one-way ANOVA; percent foot-slips of WT left hindlimb 4.8 ± 1.31%, WT right hindlimb 2.29 ± 0.47%, HRM left hindlimb 3.67 ± 0.52%, HRM right hindlimb 2.26 ± 0.64%; F _3, 28_ = 2.375, p = 0.091 one-way ANOVA). These behavioral observations that the HRM displayed impairment in skilled motor movements led us to examine the cortical map reorganization.

Cortical map reorganization is a mechanism by which mammals acquire skilled movements which emerges during the late phase of the motor training (Kleim 2004). To determine whether Reelin haploinsufficiency affects the cortical map reorganization following the motor skill training, we examined the M1 representation of the forelimb (i.e., the caudal forelimb area located in the lateral agranular field (Donoghue and Wise, 1982; Tennant et al., 2011) on the training day 10. Using intracortical microstimulation (ICMS), the forelimb movement evoked area was quantified by its rostral-medial border of the oral-facial and whisker areas, and its caudo-lateral border with the trunk and hindlimb areas (**Figure 1D**). We identified a unilateral expansion in the size of the total forelimb area, contralateral to the trained forelimb in WT trained group (**Figure 1E** n = 8-9, WT ‘ipsi’ the left cortex 0.69 ± 0.07 mm^2^, WT ‘contra’ the right cortex 0.51 ± 0.03 mm^2^, HRM ‘ipsi’ the left cortex, 0.60 ± 0.07 mm^2^, HRM ‘contra’ the right cortex 0.47 ± 0.02 mm^2^, WT trained ipsi 0.76 ± 0.06 mm^2^, WT trained contra 1.23 ± 0.13 mm^2^, HRM trained ipsi 0.83 ± 0.05 mm^2^; HRM trained contra 0.66 ± 0.07 mm^2^, F_7, 58_ = 10.10, p < 0.0001 one-way ANOVA). The cortical representation areas derived from naïve WT and naïve HRM mice were statistically indistinguishable (HRM contra vs. WT contra p > 0.999, HRM ipsi vs. WT ipsi p = 0.9891, Tukey’s multiple comparison tests). The forelimb area contralateral to the trained forelimb derived in the WT trained mice was larger in comparison with naïve WT and HRM mice (WT trained contra vs. WT contra p < 0.0001, WT trained contra vs. HRM contra p < 0.0001). In HRM mice, the forelimb area did not change in its size with training (HRM trained contra vs. HRM contra p = 0.6903). The training effect was not evident comparing the forelimb area, ipsilateral to the trained forelimb, either in WT (WT trained ipsi vs. WT ipsi p =0.9979), or in HRM (HRM trained ipsi vs. HRM ipsi p = 0.4400). The cortical thresholds used by the ICMS in evoking movements of the skeletal muscles change with cortical structural abnormalities (Nishibe et al., 2018) but not following a brain lesion (Nishibe et al., 2010) or following reach-to-grasp training (Kleim et al., 1998). We found that neither Reelin haploinsufficiency nor the reach-to-grasp training affected the cortical stimulation threshold to evoke forelimb movements by ICMS (**Figure 1F** n = 8, WT 69.1 ± 7.77 μA, HRM 61.9 ± 7.65 μA, WT trained 73.6 ± 6.67 μA, HRM trained 58.9 ± 3.16 μA; F_3. 29_ = 1.057, p = 0.382 one-way ANOVA). The amount of anesthetics mice received during the mapping procedure was ensured not significantly different among the groups (**Figure 1G** n = 8, WT 73.5 ± 5.72 mg/kg/hr, HRM 78.65 ± 9.02 mg/kg/hr, WT trained 76.14 ± 4.47 mg/kg/hr, HRM trained 84.0 ± 7.34 mg/kg/hr; F_3. 29_ = 0.430, p = 0.732 one-way ANOVA).

The anatomical connectivity of the cortico-cerebellum and cortico-striatum indicate that the cerebellum and striatum modulate cortical output (Kelly and Strick, 2003) in particular during the reach-to-grasp movements (Qian et al., 2015; Becker and Person, 2019). Binding of Reelin to its specific receptors is transduced to multiple downstream molecules via phosphorylation of Dab1 (Howell et al., 1999), suggesting Dab1 is a downstream effector of Reelin (Howell et al., 1997). To understand the importance of Reelin-Dab1 signaling in the neocortex, we examined whether motor impairment observed in the HRM also manifests in the *Dab1* ^*floxed*/ +^; *Emx1-Cre* (*Dab1* cKO) mice with intact cerebellum and striatum (Imai et al., 2017). The *Dab1* control and cKO mice were subjected to reach-to-grasp training (**Movie 3 and Movie 4**). The cKO mice showed a lower number of retrieved pellets, however, both groups improved over the training (**Figure 2A** n = 5, group effect F_1, 8_ = 6.058, p = 0.017; time effect F_2.7, 21.6_ = 3.764, p = 0.029; interaction effect F_9, 72_ = 1.04, p = 0.417, two-way repeated-measures ANOVA), similar to the behavioral results found in the HRM. Previously, the cKO mice were shown to exhibit no gross motor impairments tested on the open field and rotarod (Imai et al., 2017).

By examining the cortical forelimb map on day 10 of the training, we identified a unilateral expansion in the size of the forelimb area, contralateral to the trained forelimb (**Figure 2B, C** n = 3-4, group median (the interpercentile range of upper and lower limits) Dab1 control contra 0.342 (0.46-0.33) mm^2^, Dab1 control ipsi 0.40 (0.75-0.18) mm^2^, Dab1 cKO contra 0. 419 (0.50-0.35) mm^2^, Dab1 cKO ipsi 0.54 (0.57-0.51) mm^2^, Dab1 control trained contra 1.283 (0.35-0.27) mm^2^, Dab1 control trained ipsi 0. 56 (0.86-0.51) mm^2^, Dab1 cKO trained contra 0.921 (0.94-0.79) mm^2^, Dab1 cKO trained ipsi 0.479 (0.77-0.24) mm^2^; *χ*^2^ = 19.05, p = 0.008 Kruskal-Wallis). The training-induced expansion of the forelimb area contralateral to the trained forelimb was detected in the *Dab1* control trained mice (*Dab1* control trained contra vs. *Dab1* control contra p = 0.0458 Dunn’s multiple comparisons test) but not in *Dab1* cKO trained mice (*Dab1* cKO trained contra vs. *Dab1* cKO contra p = 0.4817). The cortical representation areas derived from naïve *Dab1* control and *Dab1* cKO mice were statistically indistinguishable (*Dab1* cKO contra vs. *Dab1* control contra p > 0.999, *Dab1* cKO ipsi vs. *Dab1* control ipsi p > 0.999). The training effect was not evident comparing the forelimb area of the M1 ipsilateral to the trained forelimb in *Dab1* control (*Dab1* control trained ipsi vs. *Dab1* control ipsi p > 0.999), or in *Dab1* cKO mice (*Dab1* cKO trained ipsi vs. *Dab1* cKO ipsi, p > 0.999). No effect of Dab1 deficiency or a training effect was observed on the cortical stimulation threshold (**Figure 2D**, n = 3-4, group median (the interpercentile range of upper and lower limits) Dab1 control 60.00 (76.6-37.0) μA, Dab1 cKO 35.71 (41.8-20.0) μA, Dab1 control trained 50.00 (50-10) μA, Dab1 cKO trained 33.00 (90-10) μA; *χ*^2^ = 1.538, p = 0.7124 Kruskal-Wallis). Neither the volume of anesthetics each mouse received was different (**Figure 2E**, n = 3-4, group median (the interpercentile range of upper and lower limits) Dab1 control 81.50 (113.5-53.3) mg/kg/hr, Dab1 cKO 83.50 (89.2-79.5) mg/kg/hr, Dab1 control trained 90.0 (93.9-47.6) mg/kg/hr, Dab1 cKO trained 76.40 (88.0-51.7) mg/kg/hr; *χ*^2^ = 2.071, p = 0.6003 Kruskal-Wallis).

The Reln haploinsufficiency or the cKO of Dab1 did not abolish the learning ability in mice but substantially reduced their level of motor skill performance. The level of behavior displayed by each group on training day 10 showed a correlation with cortical map area representing the forelimb movement (**Figure 2F**, Pearson correlation coefficient r = 0.407, p = 0.048 2-tailed). Meanwhile the level of behavior on the training day 1 showed no correlation with the forelimb motor map area (r = 0.402, p = 0.598 2-tailed). The finding is in agreement with previous rodent behavioral studies indicating that use-dependent reorganization represents the consolidation phase of training (Kleim et al., 2004). Together, Reelin haploinsufficiency affects the level of skilled motor function, and Reelin signaling may be involved in the activity-dependent plasticity of the motor map which emerges during the consolidation of motor skill.

### Altered synaptic potentiation and transmission in HRM

Cortical horizontal connections are capable of undergoing LTP which facilitates activity-dependent reorganization in M1 neurons (Hess and Donoghue, 1994; Castro-Alamancos et al., 1995). To test the hypothesis that the consequence of the genetic haploinsufficiency on the activity-dependent cortical plasticity is related to abnormality occurs earlier in the training of HRM, we evaluated the synaptic efficacy after 3 days of training, using the whole-cell patch-clamp recording. The mice used in the patch-clamp experiment exhibited a lower performance level in the pellet reach-to-grasp task similar to the HRM mice used in rest of the study (**Figure 3A,** n = 5 group effect F_1, 8_ = 7.61, p = 0.0247; time effect F_1.33, 10.72_ = 3.051, p = 0.102; interaction effect F_2, 16_ = 0.708, p = 0.507 two-way repeated-measures ANOVA). The recording was taken from the layer III pyramidal neuron which receive the sensory neuronal innervations in the motor cortex contralateral to the trained forelimb (Sievert and Neafsey, 1986). The recording from the layer III pyramidal neuron was confirmed by injecting depolarized currents to induce action potentials. To determine whether synaptic transmission undergoes LTP, synaptic stimulation was paired with postsynaptic depolarization (here referred to as paring). The paring produced a significant potentiation of synaptic responses in WT (**Figure 3B-E,** n = 12 neurons/ 5 mice t_11_ = −7.402, p < 0.001 t-test 2-tailed), HRM (n = 10 neurons/ 5 mice t_9_ = −5.293, p < 0.001), WT trained (n = 9 neurons/ 5 mice t_8_ = −4.66, p = 0.001), and HRM trained groups (n = 9 neurons/ 5 mice t_8_ = −3.707, p = 0.006). We then detected a difference in their magnitude of an increase in the synaptic potentiation (**Figure 3F**, WT 162.15 ± 8.05%, HRM 124.67 ± 5.01%, WT trained 133.40 ± 7.34%, HRM trained 119.82 ± 4.81% F_3, 36_ = 8.71, p = 0.0001 one-way ANOVA). The elicited magnitude of LTP was smaller in WT trained compared with WT group (p < 0.001, LSD), indicating that M1 synaptic potentiation had been evoked by reach-to-grasp training, which produced the occlusion of its further potentiation (Hess and Donoghue, 1994; Rioult-Pedotti et al., 1998). On the other hand, the magnitude of elicited LTP of naive HRM was reduced, and did not change with training in HRM (p = 0.185). The results show the synaptic potentiation was occluded in the basal measurements of the naïve HRM and no further training effect was achieved in the HRM, suggesting the Reelin haploinsufficiency impedes the synaptic potentiation.

Excitatory-inhibitory balance of horizontal M1 contributes to the remodeling of motor cortical representations (Jacobs and Donoghue, 1991). Thus, we examined transmissions at the excitatory and inhibitory synapses in layer III pyramidal neurons. Differential effects were detected in the frequency (**Figure 3G, H**, WT n = 18 neurons/ 5 mice 1.98 ± 0.26 Hz, HRM n = 16 neurons/ 5 mice 1.21 ± 0.18 Hz, WT trained n = 12 neurons/ 5 mice 3.20 ± 0.39 Hz, HRM trained n = 16 neurons/ 5 mice 1.68 ± 0.18 Hz; F_3, 58_ = 9.16, p = 0.0004 one-way ANOVA), and in the amplitude of the spontaneous *excitatory* postsynaptic currents (sEPSCs) (**Figure 3I**, WT 17.66 ± 0.52 pA, HRM, 17.80 ± 0.24 pA, WT trained, 21.97 ± 1.57 pA, HRM trained, 18.0 ± 0.54 pA; F_3, 58_ = 6.52, p = 0.0007 one-way ANOVA). The training increased its frequency (p = 0.002, LSD) and amplitude (p < 0.001) in WT mice, suggesting both synaptic inputs and postsynaptic function were strengthen by the training. The HRM mice exhibited a lower frequency but no difference in the amplitude compared to the WT (p = 0.032), pointing to the lower excitatory synaptic input in naïve HRM. Neither frequency (p = 0.197) nor amplitude (p = 0.850) increased by HRM training. The results suggest the *excitatory* synaptic transmission was reduced by the effect of Reelin haploinsufficiency and it was not susceptible to motor training in HRM, as it was in WT.

Furthermore, we found a significant difference in the frequency of the spontaneous *inhibitory* postsynaptic currents (sIPSCs) **(Figure 3J, K**, WT n = 17 neurons/ 5 mice 5.93 ± 0.39 Hz, HRM n = 13 neurons/ 5 mice HRM 4.22 ± 0.61 Hz, WT trained n = 13 neurons/ 5 mice 4.90 ± 0.35 Hz, HRM trained n = 17 neurons/ 5 mice 4.06 ± 0.33 Hz; F_3, 52_ = 4.14, p = 0.010) but not in the amplitude of sIPSCs (**Figure 3L**, WT 27.87 ± 0.91 pA, HRM 25.75 ± 1.49 pA, WT trained 25.71 ± 1.01 pA, HRM trained 24.84 ± 0.87 pA; F_3, 52_ = 1.56, p = 0.209). More specifically, the frequency of sIPSCs was lower in HRM than WT (p = 0.006, LSD), pointing to decreased inhibitory synaptic input in naïve HRM. The sIPSCs frequency did not change significantly in trained WT compared with WT (p = 0.092) and neither in HRM trained mice compared with HRM (p = 0.810). Hence, the *inhibitory* synaptic transmission was reduced by the effect of Reelin haploinsufficiency and it was not susceptible to motor training either in WT or in HRM mice.

Motor training on rotarod can increase or decrease the intrinsic properties of motor cortical neurons, depending on the day of the motor training (Kida et al., 2016). We investigated the possibility that Reelin haploinsufficiency impacts training-induced changes of intrinsic properties of layer III pyramidal neurons. We observed the training increased the number of spikes as injected current intensity was increased in both genotypes on the training day 3 (**Figure 4A, B**, WT n = 18 neurons/ 5 mice, HRM n = 18 neurons/ 5 mice, WT trained n = 13 neurons/ 5 mice, and HRM trained mice n = 14 neurons/ 5 mice; group effect F_3, 59_ = 4.174, p = 0.01; current intensity effect F_7, 413_ = 413.9, p < 0.0001; interaction effect F_21, 413_ = 3.420, p = 0.004 two-way repeated-measures ANOVA). The number of spikes of the WT trained was higher than that of WT (p = 0.014 LSD multiple comparison test) and HRM (p = 0.012), and the spike number of HRM trained was higher than that of WT (p = 0.014) and HRM (p = 0.02). The resting membrane potential was significantly depolarized in WT trained and in HRM trained groups compared with their respective control groups, suggesting Reelin function is probably not involved (**Figure 4C**, WT −81.03 ± 0.62 mV, HRM −80.57 ± 1.16 mV, WT trained −76.18 ± 1.01 mV, HRM trained −75.73 ± 1.21 mV; F_3, 59_ = 6.20, p = 0.0009 one-way ANOVA, WT trained vs WT p = 0.002, HRM trained vs. HRM p = 0.001, LSD). No genotype or training effect was evident in threshold (**Figure 4D**, WT −56.94 ± 1.11 mV, HRM −56.02 ± 0.60 mV, WT trained −57.40 ± 0.67 mV, HRM trained −54.87 ± 1.20 mV; F_3, 59_ = 1.27, p = 0.294 one-way ANOVA), or in input resistance (**Figure 4E** WT 143.07 ± 10.19 MW, HRM 148.73 ± 8.46 MW, WT trained 166.23 ± 11.65 MW, HRM trained 166.85 ± 12.53 MW; F_3, 59_ = 1.31, p = 0.28 one-way ANOVA). The results suggest intrinsic properties increased with training regardless of the genotype. Hence, Reelin may not be the upstream molecule in inducing changes in the neuronal membrane properties; for example, by the calcium-dependent potassium current-induced hyperpolarization (Kaczorowski and Garcia, 1999) or by TREK-1, a 2-pore-domain potassium channel currents (Honoré et al., 2006). Though no alteration in the intrinsic properties was observed in the HRM, the intrinsic properties alone did not provide a basis for synaptic modification. Together, we speculate the impairments of the synaptic plasticity detected during the early-phase of the training underlie the lack of extensive activity-dependent cortical plasticity in HRM.

**Figure 4.**
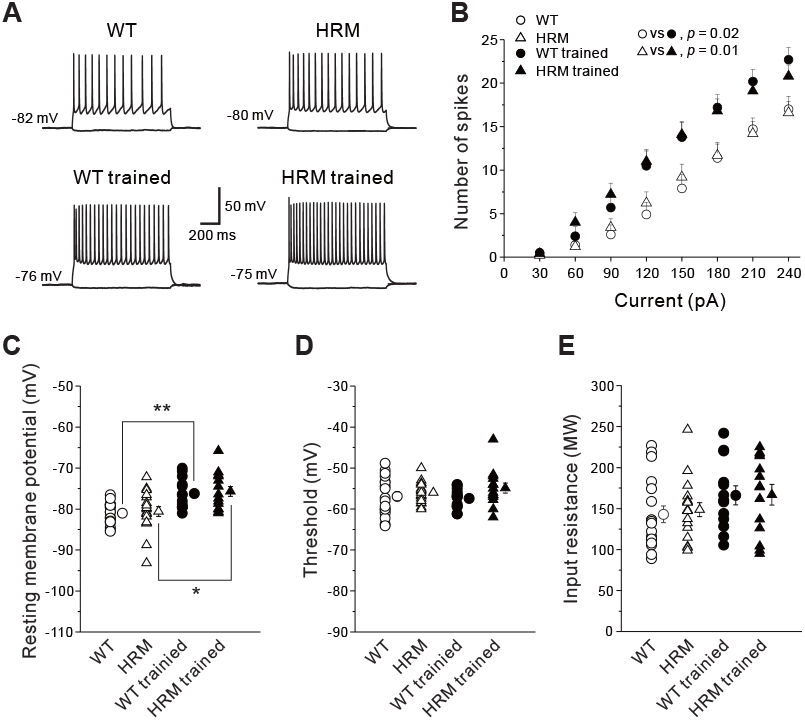
Intrinsic properties of the layer III pyramidal neurons in WT and HRM. **(A)** Representative traces of spikes obtained from cortical pyramidal neurons at respective conditions. **(B)** Relationship between current amplitude and number of evoked spikes. Data was collected from WT (n = 18 neurons/ 5 mice), HRM (n = 18 neurons/ 5 mice), WT trained (n = 13 neurons/ 5 mice), and HRM trained mice (n = 14 neurons/ 5 mice). We observed the training increased the number of spikes as the injected current intensity was increased in both WT trained and HRM trained groups. p-values of LSD post-hoc test comparing WT and WT trained; and HRM and HRM trained group are indicated. **(C)** Scatter plots and average resting membrane potential. The resting membrane potential was significantly depolarized in both WT trained and in HRM trained groups compared with their respective control groups (*p < 0.05, **p < 0.01, one-way ANOVA LSD post-hoc test), suggesting Reelin function is most likely not involved. **(D)** Scatter plots and average threshold of spikes. No genotype or training effect was evident in the threshold. **(E)** Scatter plots and average input resistance. No genotype or training effect was evident in input resistance. Data are presented as mean ± SEM.

### Altered activity-dependent gene expression in HRM

The level of mRNA synaptophysin was found reduced in the post mortem schizophrenia brain (Eastwood et al., 1995). To further assess the role of Reelin in synaptic impairments of the early training phase, we tested for expression levels of the inducible immediate-early genes, the molecules associated with LTP induction (Abraham et al., 1991; Link et al., 1995; Minatohara et al., 2015) and synaptophysin, the presynaptic marker, of the cortical M1 tissue contralateral to the trained forelimb, on the training day 3. The HRM mice exhibited a lower level of skilled motor performance (**Figure 5A**, n = 8, group effect F_1, 14_ = 6.429, p = 0.0238; time effect F_2, 28_= 1.644, p = 0.2114; interaction effect F_2, 28_ = 0.115, p = 0.8918), similar to the HRM mice used in rest of the study. The mice were then sacrificed following the completion of the day 3 of the reach-to-grasp training. Using qRT-PCR, we observed a differential expression level of Jun (**Figure 5**, primers in **Extended Data Figure 5-1** n = 8, WT 1.0 ± 0.26 relative expressions, HRM 0.71 ± 0.20, WT trained 2.07 ± 0.49, HRM trained 0.42 ± 0.11; F_3, 28_ = 5.622, p = 0.0038), Junb (WT 1.0 ± 0.25 relative expressions, HRM 0.92 ± 0.24, WT trained 2.40 ± 0.49, HRM trained 0.46 ± 0.11; F_3, 28_ = 7.462, p = 0.008) and Syp (WT 1.0 ± 0.21, HRM 0.71 ± 0.16, WT trained 2.0 ± 0.42, HRM trained 0.41 ± 0.09; F_3, 28_ = 5.069, p = 0.0063) in WT trained mice, compared to rest of the groups, but not in Arc (WT 1.0 ± 0.28 relative expressions, HRM 1.19 ± 0.41, WT trained 1.51 ± 0.37, HRM trained 0.45 ± 0.11; F_3, 28_ = 1.949, p = 0.1446), and Fos (WT 1.0 ± 0.25 relative expressions, HRM 1.05 ± 0.35, WT trained 1.72 ± 0.42, HRM trained 0.45 ± 0.10;F_3, 28_ = 2.888, p = 0.0531). The results showed that change in some of the molecular expressions associated with neuronal plasticity occurred in the WT training was not expressed in HRM, possibly contributing to the differential outcome of activity-dependent cortical plasticity.

**Figure 5.**
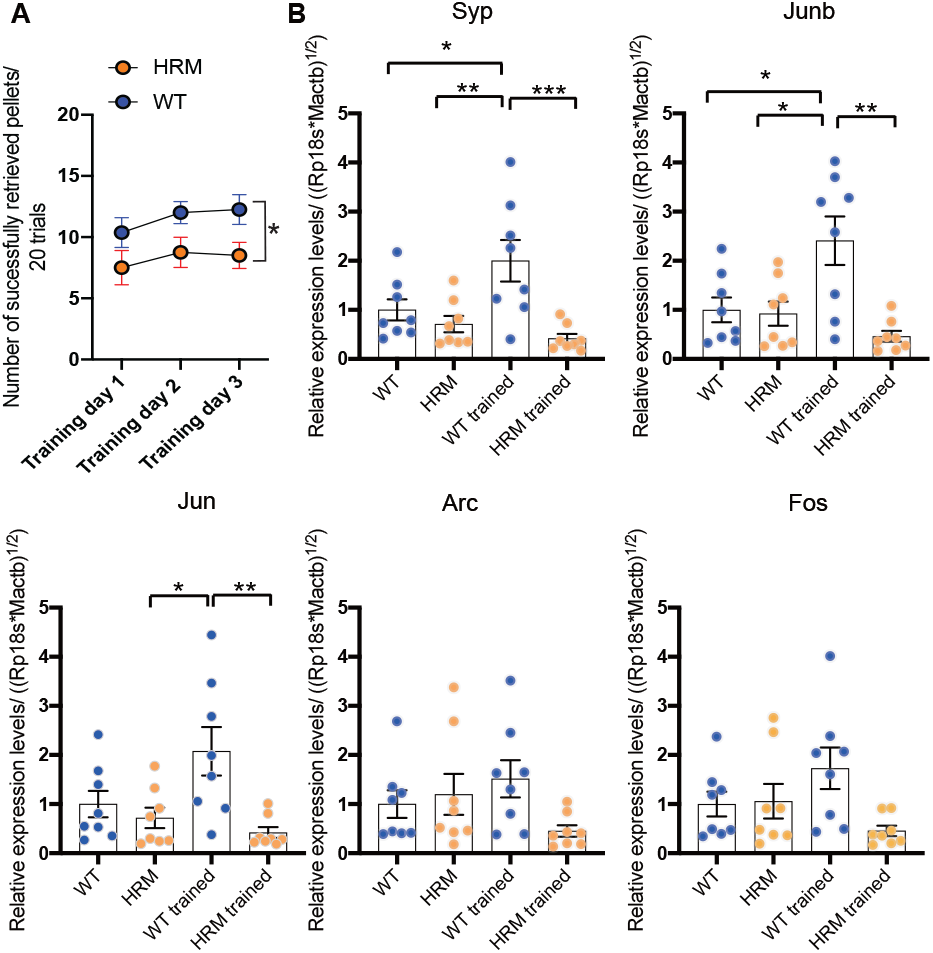
Relative expression levels of the immediate early genes and syp. **(A)** Behavior of the mice used for qRT-PCR and RNA-seq experiments. The HRM retrieved a fewer pellets than the WT (n = 8 mice, * p < 0.05 two-way repeated-measures ANOVA significant Group effect), similar to the behavior exhibited by mice used in rest of the study. **(B)** Relative expression levels of the molecular correlates of the LTP induction and presynaptic function on the training day 3, of the cortical M1 tissue contralateral to the trained forelimb (n = 8 mice); normalized by the geometric mean of the expression levels of Actb and ribosomal protein s18 (setting the value in the WT group as 1), quantified by RT-qPCR. (*p<0.05, **p<0.001, ***p < 0.001, one-way ANOVA LSD post-hoc tests). Data are presented as mean ± SEM. Primers for the genes tested are listed in Extended Data Figure 5-1.

To examine the hypothesis that Reelin plays a role in mediating certain signalings important for activity-dependent cortical plasticity, RNA-sequencing data was generated from the motor cortical tissues contralateral to the trained forelimb after 3 days of training. The analysis on the differentially expressed genes (DEGs) of the HRM group compared to the WT group (with false discovery rate cutoff of 0.1 and minimum fold change of 2 for all the following comparisons) identified no gene enrichment (**Figure 6A**), suggesting little basal difference in the transcriptome analysis in the motor cortex between WT and HRM. The DEG analysis in the WT trained group in comparison to the WT group yielded 82 upregulated and 1 downregulated genes (**Figure 6A**). The results indicate that the training exerted a greater effect on gene expression than the genetic insufficiency. The DEG analysis of the HRM trained group compared to the HRM group indicated there was no gene enrichment (**Figure 6A**), suggesting the training had little effect on the overall gene expression in HRM. Lastly, we compared the DEGs between the HRM trained and WT trained group. For this, 1 gene was upregulated and 95 genes were downregulated (**Figure 6A**). Among the DEGs, the overlap of 65 genes was found in the comparison between WT trained vs. WT, and HRM trained vs. WT trained (**Figure 6B).** Some gene validations are shown in **Figure 6C (** expression levels by RNA-sequencing n = 3 mice, Olfr50 WT −3.32 ± 0.00 log 2 expression, HRM −3.32 ± 0.00, WT trained 2.66 ± 0.19, HRM trained −3.32 ± 0.00; Olfr479 WT −2.16 ± 1.15, HRM −3.32 ± 0.00, WT trained 2.23 ± 0.41, HRM trained −3.32 ± 0.00; Olfr613 WT 9.93 ± 0.67, HRM 8.4 ± 0.57, WT trained 11.05 ± 0.32, HRM trained 7.83 ± 0.1; Gpr65 WT 1.0 ± 0.14, HRM 0.43 ± 0.05, WT trained 0.94 ± 0.15, HRM trained 0.37 ± 0.06; and expression levels by RT-qPCR n = 6 mice, Olfr50 WT 1.0 ± 0.56 relative expressions, HRM 0.12 ± 0.78, WT trained 1.45 ± 0.37, HRM trained 0.006 ± 0.004; Olfr479 WT 1.0 ± 0.44, HRM 0.11 ± 0.04, WT trained 3.36 ± 0.88, HRM trained 0.03 ± 0.008; Olfr613 WT 1.0 ± 0.34, HRM 0.50 ± 0.18, WT trained 3.36 ± 0.84, HRM trained 0.12 ± 0.051; Gpr65 WT −3.32 ± 0.00, HRM −3.32 ± 0.00, WT trained 3.32 ± 0.45, HRM trained −3.32 ± 0.00, primers are listed in **Extended Data Figure 6-1**). The upregulated pathways of both GO biological process and GO molecular function, in the comparative analysis of WT trained vs. WT were *down*regulated in the comparative analysis of HRM trained vs. WT trained (**Extended data Figure 6-2)**, suggesting the lack of gene upregulation in HRM which was induced by the motor skill training in WT. The pathway analysis indicated that G-protein coupled receptor genes, many of which were the olfactory receptor genes (Olfr), were most selectively and collectively enriched by the motor training. The results indicate that the effect of genetic insufficiency becomes apparent only when mice were intervened behaviorally; and that Reelin is at least partially involved during the early phase of the cortex-dependent motor skill training at the level of transcriptome.

**Figure 6.**
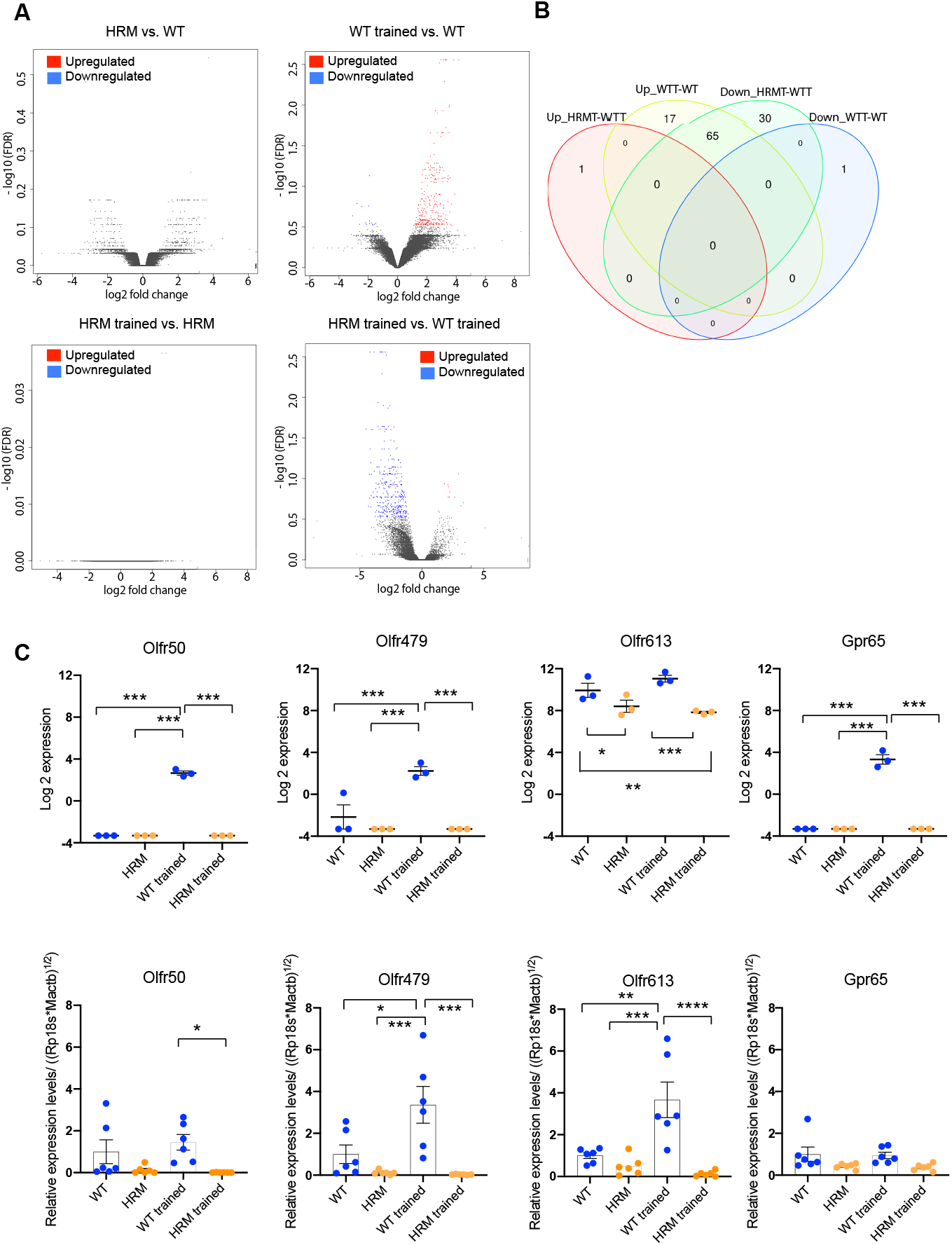
Gene upregulations in ‘trained’ M1 after reach-to-grasp training in WT and HRM. **(A)** Volcano plots of (TOP) HRM vs. WT, WT trained vs. WT, (BOTTOM) HRM trained vs. HRM, and HRM trained vs. WT trained groups comparative analyses of the RNA-seq (n = 3 mice). **(B)** Venn diagram indicating the number of upregulated and downregulated genes, identified by the RNA-seq DEG analysis of WT trained vs. WT, and HRM trained vs. WT trained. The FDR cutoff was set at 0.10 and minimum fold change at 2 for all comparisons. **(C)** Validation of some gene expressions implicated by the RNA-seq. TOP: gene expression levels of Olfactory receptor (Olfr) 50, Olfr479, Olfr613, and G-protein coupled receptor 65 (Gpr65), measured by RNA-sequencing. (n = 3 mice, *p < 0.05, **p < 0.01, ***p < 0.001, one-way ANOVA LSD post-hoc tests). BOTTOM: validation of these gene expressions by qRT-PCR, normalized by the geometric mean of the expression levels of Actb and ribosomal protein s18 (setting the value in the WT group as 1). (n = 6 mice, *p < 0.05, **p < 0.01, one-way ANOVA LSD post-hoc tests). Data are presented as mean ± SEM. Primer are listed in Extended Data Figure 6-1

## Discussion

Deterioration of Reelin mRNA level is indicated in the brain of those with psychiatric disorders (Impagnatiello et al., 1998; Guidotti et al., 2000). Investigations of genome sequence identified mutations in *RELN* and *DAB1* genes in patients with psychiatric symptoms (Ovadia and Shifman, 2011; Teroganova et al., 2016). Thus, the molecular features exhibited by HRM at least partially resemble some features observed in psychiatric disorders. Our primary findings are that mice with Reelin haploinsufficiency exhibited lower cortex-dependent skilled motor performance; and that the effect of genotype emerged in the gene expression, synaptic plasticity and cortical map plasticity in response to the motor skill training.

When subjected to the single pellet reach-to-grasp training, HRM and *Dab1* cKO mice did not show a loss in the ability to learn though there was no substantial map reorganization. It is possible other motor areas including the cerebellum and striatum supported their increase in the motor skill performance; particularly these structures of *Dab1* cKO mice were assumed normal. Nevertheless, disassociation of motor *learning* and map reorganization has been observed in aged mice, or during recovery from a brain injury. In aged mice, the ability to learn a skilled motor task is maintained with a lack of extensive cortical map reorganization (Tennant et al., 2012). The map reorganization and motor behavior are dissociated during sub-acute phase of a brain injury (Nishibe et al., 2015). In the present study, the degree of map reorganization as a result of competitive map interaction was associated with the level of motor performance during the late phase of the training and not during the early phase of the training, in agreement with a previous rodent study, suggesting the map plasticity represents the consolidation of motor skill that occurs during the late training stages and not the acquisition during the early stages (Kleim et al., 2004). During the early phase of the training, dendritic and synaptic hypertrophy (Greenough et al., 1985) and dendritic pruning (Xu et al., 2009) are shown to occur, but the formation of new synapses is detectable only after motor skill acquisition (Kleim et al., 2004). Our results indicate that the excitatory synaptic transmissions, synaptic potentiation and intrinsic properties were strengthened as well as certain genes were enriched, potentially contributing to the initial acquisition of motor skill during the early phase, and subsequently constructing the late-phase activity-dependent cortical plasticity.

We examined the activity-dependent cortical plasticity of the caudal forelimb area in the lateral agranular field that has shown to undergo redistribution of body representations (Kleim et al., 1998); apart from the rostral forelimb area in the medial agranular field (Donoghue and Wise, 1982) where the area receives less sensory inputs than the caudal forelimb area (Sievert and Neafsey, 1986). The cortical reorganization of the caudal forelimb area appeared to be mediated by the excitatory synapses in WT, through an increase in the spontaneous action potential driven glutamatergic transmission of both strengthened presynaptic and postsynaptic effects, providing a putative mechanism of LTP. Also, increases in the expression level of immediate early genes provide a molecular basis in inducing LTP and cortical reorganization possibly at least partially through Reelin signaling. Reelin regulates synaptic formation by changing the molecular composition of the synapses (Sinagra et al., 2005), or by regulating the stability of dendritic spines (Niu et al., 2004; Ventruti et al., 2011). In the brain of HRM, synaptic structural abnormalities have been reported, including decreased number of spines and reduced level of glutamic acid decarboxylase 67 expressions (Liu et al., 2001; Qiu et al., 2006; Hellwig et al., 2011; Bosch et al., 2016). Consistent with the previous observations, we found that the basal LTP, excitatory and inhibitory synaptic inputs of M1 layer III neurons were decreased in naïve HRM compared with naïve WT. The observation implies that there is a decrease in the presynaptic excitatory release and decrease in the levels of postsynaptic glutamate receptor. Also, the result reflects the presence of abnormal spine formations and declined number of spines in HRM. The existing occlusion of LTP and depressed synaptic inputs in naïve HRM did not compromise the overall integrity of the basal cortical map; however, impeded the occurrence of further horizontal LTP and use-dependent map reorganization. Although we detected an increase in the neuronal resting membrane potential both in the WT and HRM group following the training; such increase alone did not contribute to the LTP-induction or substantial map reorganization in HRM. Hence, in line with the previous reports (Rioult-Pedotti et al., 1998; Rioult-Pedotti et al., 2000), extensive synaptic modification is required for the development of extensive map reorganization. The finding reveals the synaptic impairments that were apparent in HRM mice suppressed the neuronal adaptation imposed by the motor skill training. Thus, the impairment of the activity-dependent cortical plasticity is speculated to be closely linked to the depressed synaptic function whether it is caused developmentally or by *de novo* abnormalities.

To further demonstrate the effect of genetic insufficiency during the early phase of the training, we showed the absence of differential expressions of some of the tested immediate early genes (IEGs) associated with the induction of LTP, and synaptophysin in HRM trained mice. Whether these IEGs that were not explicitly expressed in the HRM trained mice in the same extent in the WT trained mice are under the direct or indirect control of Reelin requires further investigation. Using whole genome analysis, we provided compelling evidence that the response of gene expression to the motor experience requires normal level of Reelin signal activity. It appears that the training effect was larger in the outcome of cortical gene expressions than the effect of genetic insufficiency. This was also true in the activity-dependent cortical map representation, similar to a clinical finding in patients with brain-derived neurotrophic factor (BDNF) polymorphism (Kleim et al., 2006), implicating that the effect of genetic insufficiency became apparent only when subjects were intervened behaviorally.

It is worth considering that Reelin regulates molecular pathways indicated by RNA-sequencing directly or indirectly in an activity-dependent manner. Unexpectedly, our pathway analysis suggested that G-protein coupled receptor (GPR) genes, many of which were the Olfr, the largest group of the gene family in the genome, were the group that was most selectively and collectively enriched by the motor training. A number of “ectopic” Olfrs are expressed in the cerebral cortex of mice (Otaki et al., 2004). In cultured dopaminergic neurons expressing Olfr genes, odor-like agonist can trigger intracellular calcium increase (Grison et al., 2014). Our result on the Olfr expression levels in relation to the activity-dependent cortical plasticity requires further investigation. Nonetheless, the reduction of Olfr genes has been demonstrated in the frontal cortex of patients with Alzheimer’s disease (Ansoleaga et al., 2013), schizophrenia (Ansoleaga et al., 2015), and Parkinson’s disease (Garcia-Esparcia et al., 2013) for which pathological associations with deteriorated Reelin signal is documented. Also, PIK3CA-AKT1 signaling, the cascade activated by Reelin (Beffert et al., 2002; Bock et al., 2004), specifically mediated by GPR is implicated in Parkinson’s disease (Nakano et al., 2017). The present study would support for further studies on the contribution of Reelin-mediated mechanisms including receptor insertion and intracellular signaling.

Collectively, the present study demonstrated that at least in part Reelin signaling plays a role in the cortical plasticity that supports the development of skilled movement. The effect of genotype was found in the suppression of the gene enrichment, synaptic function and activity-dependent cortical map plasticity. We propose the lack of gene enrichment and synaptic abnormality led by the Reelin haploinsufficiency, which were apparent during the early phase of the training, potentially contribute to impede the development of extensive activity-dependent cortical plasticity.

## Acknowledgments

We sincerely thank Dr. Daisuke Okuzaki for the RNA-seq processing on the samples submitted to the Genome Information Research Center at Osaka University. This work was funded by the Japan Society for the Promotion of Science KAKENHI (19K11366) and Research Grants from The Nakatomi Foundation to M.N; KAKENHI (18K06502) to Y.K.; and KAKENHI (17K08538) to H.T.

---

**Extended Data Figure 5-1:**
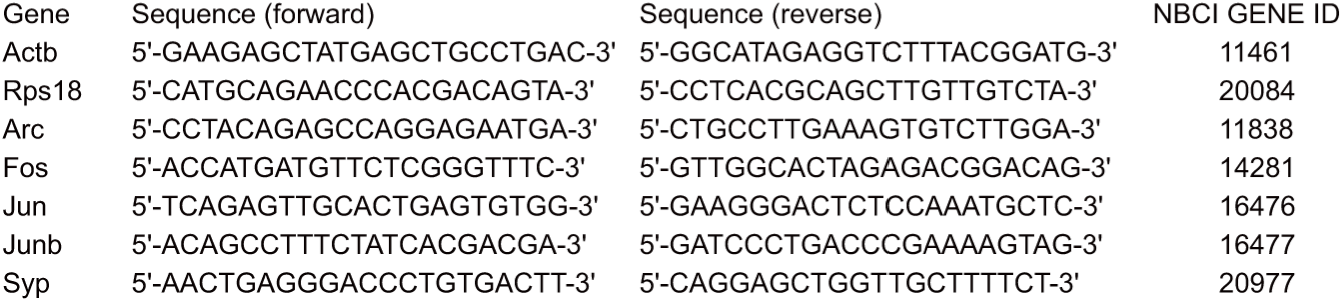
the list of primers used in the qRT-PCR.

**Extended Data Figure 6-1:**
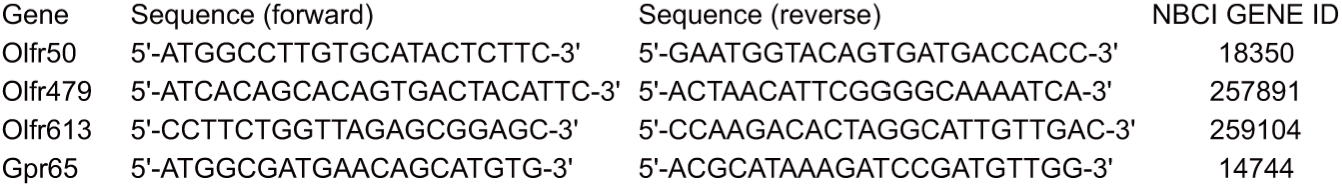
the list of primers used in the qRT-PCR.

**Extended Data Figure 6-2:**
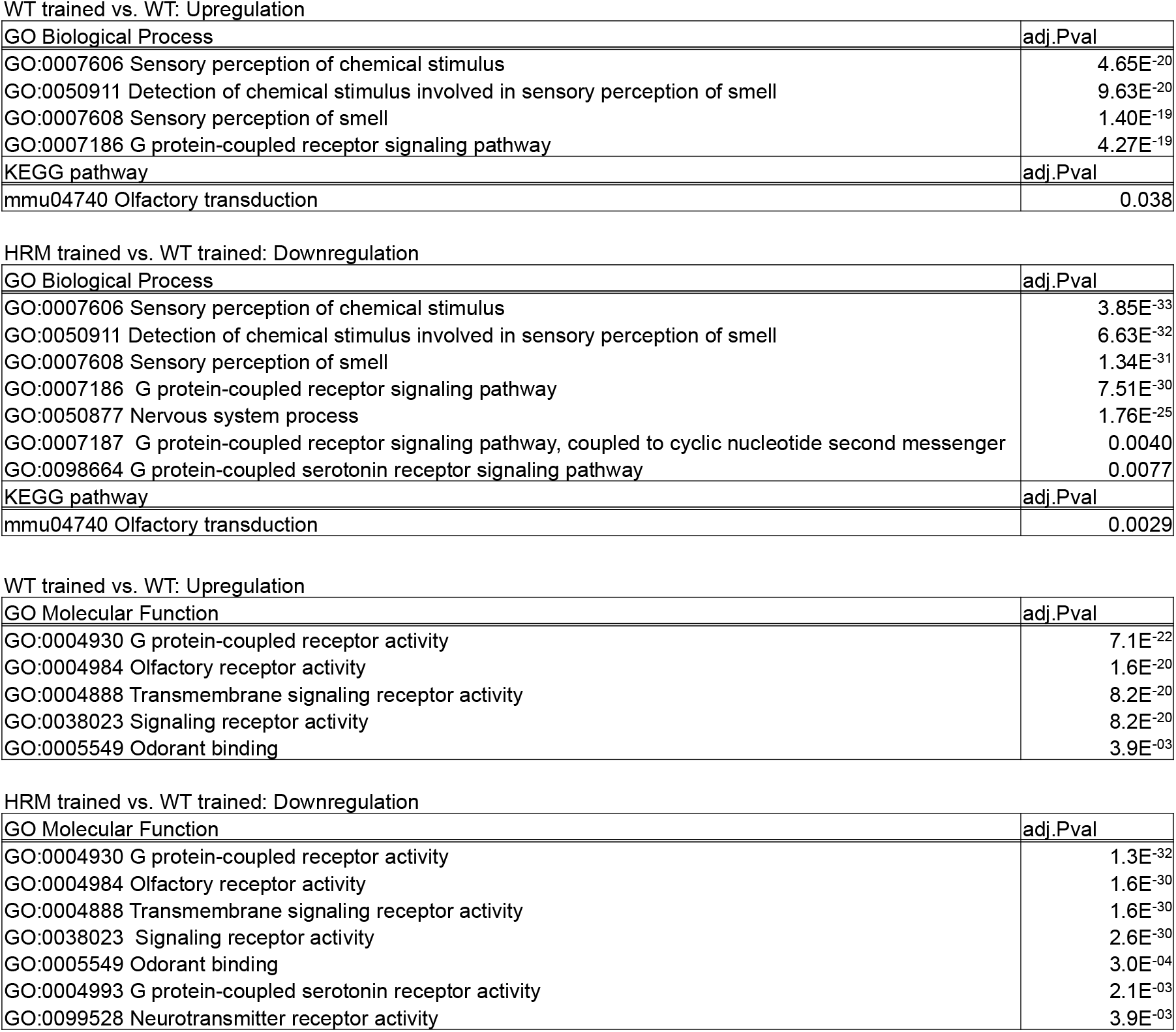
enriched GO pathways of biological process and molecular function, of WT trained vs. WT, and HRM trained vs. WT trained analysis. TOP: list of upregulated gene ontology (GO) biological process pathways and KEGG pathway in the comparative analysis of WT trained vs. WT, reported with pathway ID and each adjusted p value; followed by list of downregulated GO biological process pathways and KEGG pathway in the comparative analysis of HRM trained vs. WT trained, reported with pathway ID and each adjusted p value. BOTTOM: list of upregulated gene ontology (GO) molecular function pathways in the analysis of WT trained vs. WT, reported with pathway identification number and each adjusted p value; followed by list of downregulated GO molecular function pathways in the analysis of HRM trained vs. WT trained, reported with pathway identification number and each adjusted p value. The FDR cutoff was set at 0.10 and minimum fold change at 2 for all comparisons.

**Movie 1.** Training of the WT mouse at the single pellet reach to grasp task. This video was taken on the day 5 of the training.

**Movie 2.** Training of the HRM mouse at the single pellet reach to grasp task. This video was taken on the day 5 of the training.

**Movie 3.** Training of the *Dab1* control mouse at the single pellet reach to grasp task. This video was taken on the day 5 of the training.

**Movie 4.** Training of the *Dab1* cKO mouse at the single pellet reach to grasp task. This video was taken on the day 5 of the training.

## Notes

**Conflict of interest statement:** All authors declare no competing interests.

### Competing Interest Statement

The authors have declared no competing interest.

